# Modulating brain networks with transcranial magnetic stimulation over the primary motor cortex: a concurrent TMS/fMRI study

**DOI:** 10.1101/2019.12.16.878439

**Authors:** JeYoung Jung, Andreas Bungert, Richard Bowtell, Stephen R. Jackson

## Abstract

Stimulating the primary motor cortex (M1) using transcranial magnetic stimulation (TMS) causes unique multisensory experience such as the targeted muscle activity, afferent/reafferent sensory feedback, tactile sensation over the scalp and ‘click’ sound. Although the human M1 has been intensively investigated using TMS, the experience of the M1 stimulation has not been elucidated at the whole brain. Here, using concurrent TMS/fMRI, we investigated the acute effect of the M1 stimulation of functional brain networks during task and at rest. A short 1Hz train of TMS pulses applied to individuals’ hand area in the M1 during motor execution or at rest. Employing the independent component analysis (ICA), we showed the M1 stimulation decreased the motor networks activity when the networks were engaged in the task and increased the deactivation of networks when the networks were not involved in the ongoing task. The M1 stimulation induced the activation in the key networks involved in bodily self-consciousness (BSC) including the insular and rolandic operculum systems regardless of states. The degree of activation in these networks was highest at rest compared to during task, showing the state-dependent TMS effect. Furthermore, we demonstrated that the M1 stimulation modulated other domain general networks such as the default mode network and attention network and the inter-network connectivity between these networks. Our results showed that the M1 stimulation induced the widespread changes in the brain at the targeted system as well as non-motor, remote brain networks, specifically related to the BSC. Our findings shed light on understanding the neural mechanism of the complex and multisensory experience of the M1 stimulation.

## Introduction

TMS is a non-invasive brain stimulation technique that has been widely used to investigate brain function. TMS induces a brief magnetic field at the scalp which can then transiently modulate brain electrical activity leading to behavioural changes (Walsh and Cowey, 2000). In human brain, the primary motor cortex (M1) has been intensively investigated using TMS to understand the neural mechanism of motor system (Wasserman et al., 2008). Different from other regions, the M1 stimulation elicits motor-evoked potential (MEP), a measurement of corticospinal excitability, accompanying with a unique sensation at the targeted hand region or hand twitch at suprathereshold intensity. Studies on the M1 stimulation has focused on the MEP linking to neurophysiological mechanism of motor system but the whole experience of M1 stimulation (e.g., afferent/reafferent sensory feedback, tactile sensation over the scalp, and ‘click’ sound) has been disregarded. Thus, it is lacking in understanding how the experience of M1 TMS influences the neural processing.

The M1 TMS affects both the targeted area and functionally connected remote areas (Ferbert et al., 1992;Di Lazzaro et al., 1999;Koch et al., 2006). To demonstrate it, there have been a number of attempts to combine TMS with other neuroimaging techniques (e.g., fMRI). Bohning firstly demonstrated that TMS induced blood-oxygenation-level-dependent (BOLD) changes at the target site and a number of remote areas when TMS was applied on the M1 (Bohning et al., 1998). Since then, studies have used this technique to demonstrate that TMS can induce widespread alterations in the brain. Since the first successful demonstration of the combination of TMS and fMRI, several studies have been published using concurrent TMS/fMRI (Bohning et al., 1999;2000a;2000b;Baudewig et al., 2001;Bestmann et al., 2003;Bohning et al., 2003;Bestmann et al., 2004;Li et al., 2004;Bestmann et al., 2005;Denslow et al., 2005;2008;Ruff et al., 2009). Initial studies to stimulate M1 at rest demonstrated that TMS caused significant BOLD changes in the target region, functionally connected cortical and subcortical motor regions as well as non-motor regions such as auditory cortex, insular, frontal and parietal regions (Bohning et al., 1999;2000a;2000b;Bestmann et al., 2003;2004;Denslow et al., 2005). Recently, Bestmann et al (2008) stimulated the left premotor cortex (PMC) during a motor execution task (grip vs. no-grip) and demonstrated state-dependent TMS effect in the contralateral M1 and PMC. These findings suggest that TMS induces neural changes across the whole brain covering the target region and other remote areas and the current state of a targeted neural system influences the effect of TMS. However, studies have focused on the targeted network – the motor system at the regional activity level, disregarding other multisensory processing evoked by the M1 TMS. In addition, it remains unclear how TMS over the M1 modulates the brain at a network level covering the targeted network and other functional neural systems, depending on the current state of the network.

Here, we examined the acute effect of the M1 stimulation in brain networks during two different states (task or rest) using the concurrent TMS/fMRI. We applied a short burst of M1 stimulation during a motor execution task or at rest. A group of participants performed either unimanual or bimanual hand clenching with the left or right M1 stimulation. The other group received the left M1 stimulation without a task. As a control group, there was the vertex stimulation group at rest. To delineate brain networks from fMRI data, we employed independent component analysis (ICA) – a data-driven multivariate approach to decompose a mixed signal into independent components (networks) (Calhoun et al., 2001). We hypothesized that the ICA would identify several networks related to M1 stimulation as well as task and reveal experimental condition-specific modulation at each networks.

## Materials and methods

### Participants

Thirty-six healthy subjects (9 males: mean age 26 ± 5 years, range 19-32 years) were recruited for this study. Subjects were allocated into three groups: task M1 stimulation group, rest M1 stimulation group and rest vertex stimulation group as a control group. The task M1 stimulation group (twelve healthy adults, 3 males: mean age 27 ± 3 years, range 20-32 years) performed a motor execution task with the M1 stimulation. The rest M1 stimulation group (twelve healthy adults, 3 males; mean age 25◻±◻3.1◻years, range 19–30◻years) received the M1 stimulation without a task. As a control group, we had the vertex stimulation group from previously published study (Jung et al., 2016) (twelve healthy adults, 3 males; mean age 25◻±◻8.3◻years, range 20–28◻years) received the vertex stimulation at rest. They were all right-handed. Handedness was assessed by the Edinburgh handedness inventory (mean score: 95 ± 12) (Oldfield, 1971). All participants provided an informed written consent in advance the experiment. This study was approved by the local ethics committee and performed in accordance with the Declaration of Helsinki.

### Experiment design and Procedure

We used an fMRI block design with a block length of 30s. During the task M1 stimulation, there were four sessions in the experiment and each session comprised of 9 blocks (4mins 30s) (Fig. 1). In each session, there were four experimental conditions. Participants were asked to perform simple hand clenching movements using their left (LHC), right (RHC), or both hands simultaneously (BHC), or else they were instructed to make no hand movements (rest). The hand clenching task required participants to continuously clench and unclench their hand while instruction words were displayed on the screen (left, right, both and rest). The order of task conditions was pseudorandomized and counterbalanced across the session. Each block was comprised of a TMS phase (11s) and a No-TMS phase (19s). The temporal onset of the TMS phase was randomized within each block. During the TMS phase, 12 pulses of 1 Hz TMS were delivered to the hand area of the left or right M1. Two sessions involved TMS delivered to the left M1 and two sessions involved TMS delivered to the right M1. After the first two sessions, the TMS coil was re-positioned for the contralateral M1. The order of left and right M1 stimulation was counterbalanced across participants. The same fMRI paradigm with 18 blocks (9mins) was used for the rest M1 stimulation and the vertex stimulation without task.

**Fig. 1.**
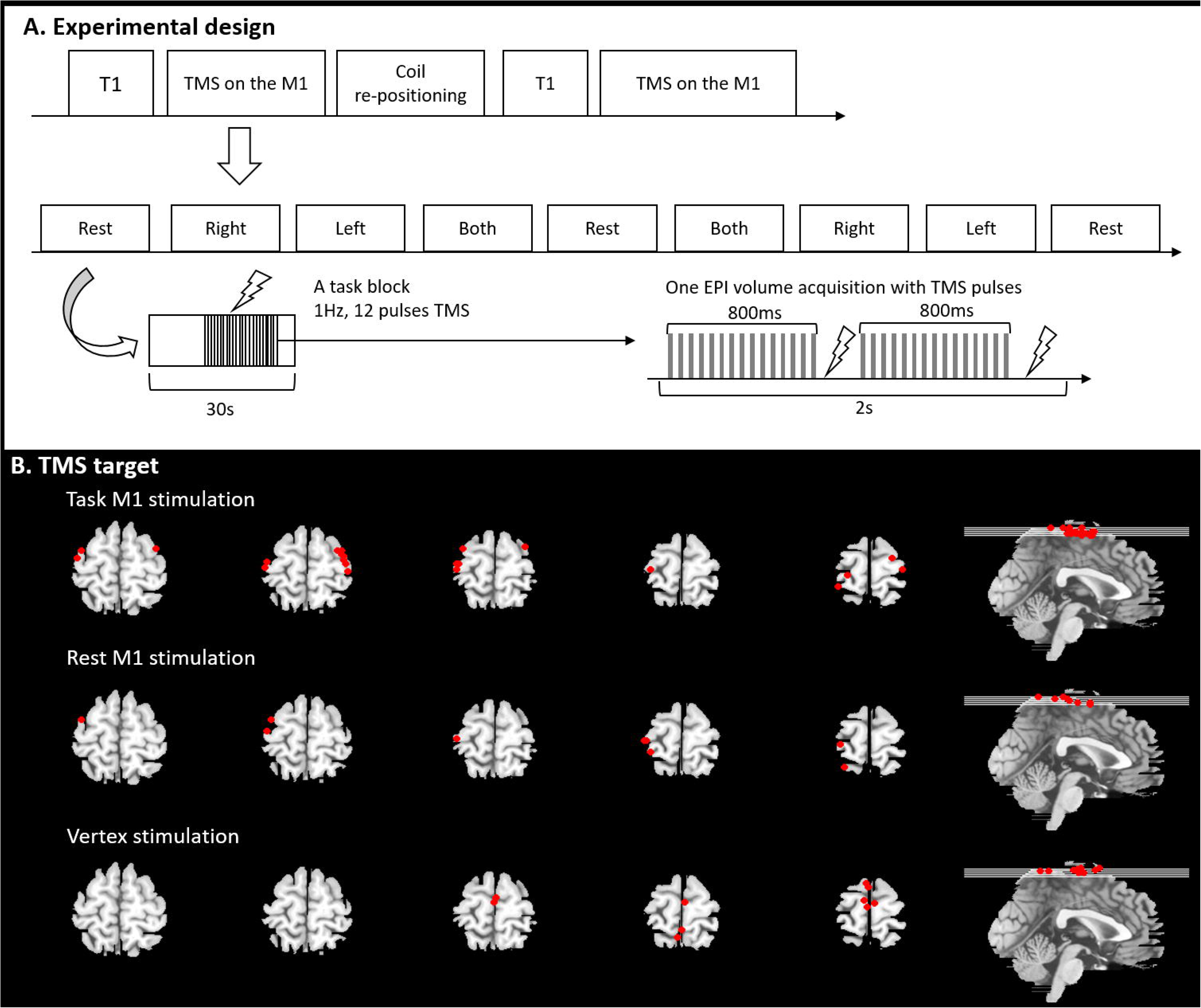
A) Experimental design and procedure. B) TMS target position.

Before the experiment, individual resting motor threshold (RMT) was measured for all participants outside of MRI scanner. Once placed in the MR scanner, the participants’ hands were placed next to their body in a natural relaxed position. Each participant wore glasses with a prismatic mirror to view a projection screen on which visual stimuli were presented throughout the experiment. The task M1 stimulation group was asked to perform the hand clenching task at a rate that they were comfortable with. Prior testing confirms that this is between 0.5-1Hz. The instruction words for each condition were displayed on a projection screen at the foot of the scanner bed. The experimental design and procedure is illustrated in Fig 1A. The other groups were instructed to keep their limbs relaxed during the experiment.

### Magnetic resonance imaging

Functional MR images were acquired at the Magnetic Resonance Centre (University of Nottingham), using a Philips 3.0-Tesla scanner equipped with a 6-channel head coil to accommodate the TMS coil. Functional images were obtained using single-shot echo planar imaging (EPI) sequence (repetition time (TR)/echo time (TE) = 2000/35 ms, flip angle 90°, 30 slices, matrix = 64 × 64, 3 × 3 × 3mm^3^ resolution). Anatomical images were acquired using 3D MP-RAGE sequence (TR/TE = 8.278/2.3 ms, flip angle 8°, matrix = 192 ×192, 1 × 1 × 1mm^3^ resolution) covering the whole head. During scanning all participants wore ear-plugs with head cushioned by foam pads to prevent head-movement artifacts.

### Transcranial magnetic stimulation

A Magstim Rapid2 stimulator (Magstim, UK) was used to generate TMS pulses through a MR-compatible figure-of-eight coil (70mm outer wing diameter). For the M1 stimulation, the coil was centred at the left or right hand area. The coil was positioned at the vertex (Cz) using the international 10-20 system (Steinmetz et al., 1989) for the vertex stimulation. Individual RMTs were measured as follows. TMS pulses were applied to M1 to identify the optimal site eliciting a muscle twitch in the left and the right FDI muscle and was oriented perpendicular to the central sulcus at a 45° angle from the mid-sagittal line approximately. Once a site was identified, the stimulator intensity was systematically varied and RMT was defined as the minimum stimulator output that was required to induce an observable muscle twitch at that site for five out of ten TMS pulses. Individual TMS intensity was 100% of RMT for the M1 stimulation. The vertex stimulation group received TMS pulses with 120% of RMT. The mean stimulator output corresponding to RMT was 75% for the right M1 stimulation (range 59 - 88%) and 75% for the left M1 stimulation (range 64 – 89 %). In the rest M1 stimulation, the mean RMT was 72% ranging from 59% to 86%. The mean RMT of the vertex stimulation was 72% ranging from 59% to 86%. To hold and fix the TMS coil, we used a plastic coil holder placed next to the MRI head coil.

### Synchronisation TMS and fMRI

Our previous study (Jung et al., 2016) showed a successful synchronization TMS and fMRI based on the findings of Shastri et al (1999). In this experiment, the scanner sequence was programmed to split the acquisition of images in each volume into two separate packages. The first package was acquired for ~800ms and the second package commenced collection 200ms after the first package acquisition had ceased. We applied a TMS pulse 850ms after the acquisition of the first slice in each package during the TMS phase of each block. In this way, TMS was applied at 850ms and at 1850ms during each volume acquisition without distortion (Fig. 1A). The synchronisation of TMS pulse was carried out using an in-house Matlab programme written in Matlab (R2006b).

### TMS coil position and target site

The rubber ring around the TMS-coil is MR-visible for short echo-time (TE<10ms) and it can be used to verify the position of the TMS-coil relative to the subject. For this purpose magnetisation prepared rapid gradient echo (MP-RAGE) images were acquired for each position of the TMS-coil. These images covered the head of the subject and the rubber ring around the coil. Using in house-Matlab code, several points on the rubber ring were identified in the images. By fitting the shape of the rubber ring around the coil to these points in the image, the position of the TMS were determined. The location defined as coil position, was the point were a virtual line perpendicular to the TMS-coil and through the centre point of the TMS coil (where the two rings of the figure-of-eight meet each other) hits the brain surface. This position was translated into MNI space by coregistering the MP-RAGE image to MNI space. TMS target sites were displayed in Fig 1B and Table S1.

### Univariate analysis

Statistical Parametric Mapping software (SPM12, Wellcome Department of Imaging Neuroscience, UK) was used for data analysis. All EPI images were re-aligned, co-registered with each individual’s anatomical image, spatially normalised to Montreal Neurological Institute (MNI) space, and spatially smoothed using a Gaussian kernel (8mm, Full-width half-maximal). The session with head movements exceeding more than 2mm in x, y, z direction and 2° in rotation, was excluded from the analysis. Head movement parameters were included within the analysis as regressor variables to exclude head movement related variance.

A general linear model (GLM) was used to calculate individual contrasts. For the task M1 stimulation group, we defined a design matrix comprising task conditions (LHC, RHC, BHC, and rest) and TMS phases (TMS and noTMS). T-contrasts for each condition and TMS phase were established for all participants. In the group analysis, three-factorial ANOVA with the site of stimulation (left vs. right), task (LHC, RHC, and BHC), and TMS (TMS vs. noTMS) was conducted and contrasts were entered into a set of one-sample *t* tests for the three movement conditions and TMS phases. For the rest M1 stimulation and vertex stimulation groups, a design matrix with TMS phases (TMS and noTMS) was constructed. In the group level analysis, the contrast images were entered into one-sample *t* tests. Statistical significance threshold was set to a height threshold of p < 0.005 uncorrected, at the voxel level and to that of p < 0.05 at the cluster level with at least 20 contiguous voxels after false-discovery rate (FDR) correction.

To explore the effect of TMS within the motor system, regions of interest (ROIs) were defined as a 4mm radius sphere based on the results of group-level analysis in the task M1 stimulation group. These included the M1 (Ml [−33, −24, 63], Mr [36, −15, 54]), PMC (PMCl [−50, −15, 37], PMCr [52, −12, 38]) and supplementary motor areas (SMA: SMAl [−3, −6, 48], SMAr [6, −6, 48]) in both hemispheres. ANOVAs with the site of stimulation (left vs. right), TMS (TMS vs. NoTMS), and hemisphere (left vs. right) as within subject factors were conducted for each ROI, according to tasks (LHC, RHC and BHC). Also, we performed a conjunction analysis to assess the general effect of TMS across the task conditions.

### Multivariate analysis – Independent component analysis

Independent component analysis (ICA) was used to estimate spatiotemporal functional networks from the data. ICA uses fluctuations in the fMRI data to separate the signal into maximally independent spatial maps or components, each explaining unique variance of the 4D fMRI data. Each component has a time course related to a coherent neural signal potentially associated with intrinsic brain networks, artefacts, or both.

ICA was performed using the group ICA of fMRI Toolbox (GIFT) (Egolf et al., 2004). The pre-processed data was entered into the GIFT version 3.0b. The toolbox concatenates the individual data followed by the computation of subject-specific components and time course. Maximum Description Length (MDL) and Akaike’s criteria were applied to estimate the number of independent components (ICs) in our data. Using principal component analysis, individual data was reduced. Then informax algorithm (Bell and Sejnowski, 1995) was applied for the group ICA and estimated 14 components. In order to improve the IC’s stability, ICASSO was applied and run 20 times (Himberg et al., 2004). Of 14 components, one component related to residual artefact was excluded for further analysis.

13 components were correlated with the brain and defined as brain networks. We labelled them with regional or functional descriptors (e.g., default mode network; DMN, motor network; MN). Then, we examined the experimental condition-relatedness in each network using the temporal sorting in GIFT. Temporal sorting applied the GLM to the component’s time course. The fMRI specific time course for each individual were regressed against the design matrix for the experimental conditions and tested for significance to identify networks where activity was greater during each condition. The resulting β weights represent the degree to which network was recruited by the conditions. For a network, positive and negative β weights indicate the level of network recruitment in each experimental condition. To assess it, one-sample *t* tests were conducted on β weights (p _FDR-corrected_ < 0.05) in each group. Then, the networks showed the significant recruitment in the experimental conditions were selected for the next analysis. For the task M1 stimulation group, a two-factorial ANOVA with the site of stimulation (left and right) and TMS (TMS and noTMS) as within-subject factors was performed for the left M1 stimulation and the right M1 stimulation according to each task (LHC, RHC, BHC and rest) separately. For the comparison between TMS and noTMS phase, *post hoc* paired *t* tests were performed for each network (p < 0.05). In order to evaluate the effect of the M1 stimulation during task and at rest relative to the control stimulation, one-way ANOVA was conducted across the groups on the resting condition. *Post hoc t* tests were performed for each network to compare the effect of TMS site (M1 and vertex) (p < 0.05).

To assess the connectivity between networks, functional network connectivity (FNC) analysis was performed. The FNC was estimated as the Pearson’s correlation coefficients between pairs of time-courses of networks (Jafri et al., 2008). To explore the FNC changes caused by the group (task M1 [left M1 TMS], rest M1 and vertex stimulation), ANOVA was conducted and following *post hoc* two-sample *t* tests were performed (p < 0.05).

## Results

### GLM results

The GLM results are summarised in Fig. 2A and Table S2. The main effect of task revealed that hand movements increased brain activity in the motor system including the bilateral M1, PMC, and SMA as well as primary sensory cortex (S1), putamen, thalamus and visual cortex. The main effect of TMS was found in the bilateral superior parietal lobe (SPL), precuneus, middle cingulate cortex (MCC) and left superior temporal gyrus (STG). The main effect of TMS site showed the activation in the bilateral M1/S1, inferior frontal gyrus (IFG), left insular, right putamen, STG and middle temporal gyrus (MTG). There was a significant interaction between the site and task in the bilateral M1 and S1. No voxels were survived in the other contrast of interactions.

**Fig. 2.**
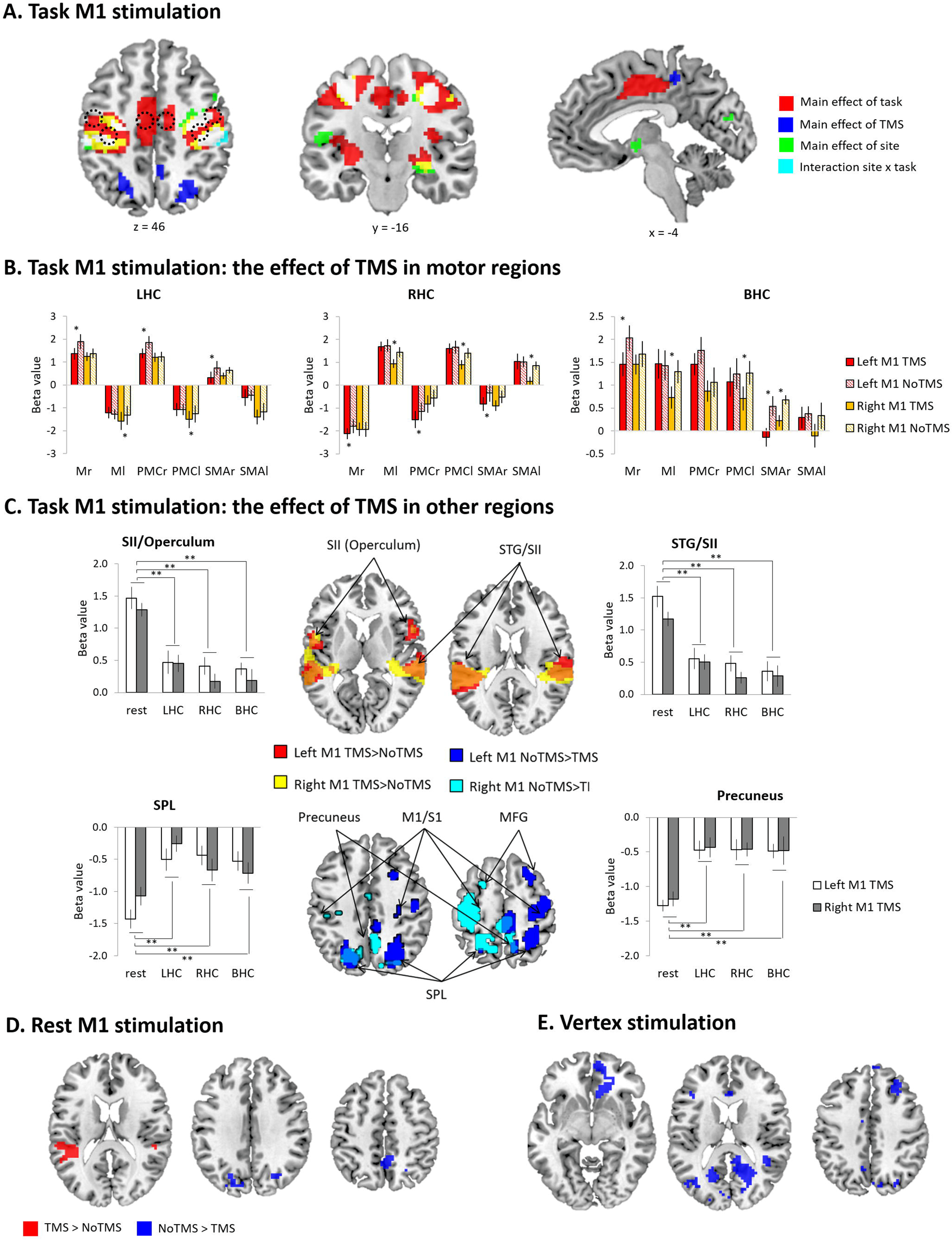
GLM results A) task M1 stimulation whole brain activation map. Red (main effect of task), Blue (main effect of TMS), Green (main effect of site), Cyan (interaction between task and site), and White (overlapping areas across the contrasts) B) The results of ROI analysis in the motor regions (M1, PMC, and SMA): Task M1 stimulation. The bar filled colour indicates the TMS phase and the bar filled with diagonal lines indicates the No TMS phase. Red (left M1 TMS) and Yellow (right M1 TMS). C) The results of conjunction analysis: Task M1 stimulation. D) Rest M1 stimulation whole brain activation map. Red (TMS > NoTMS) and Blue (NoTMS > TMS). E) Vertex stimulation whole brain activation map. Red (TMS > NoTMS) and Blue (NoTMS > TMS). Error bars indicate standard errors. * p < 0.05

To investigate the task-specific TMS effects within the MN, repeated-measures ANOVAs with hemisphere (left vs. right), site of stimulation (left vs. right) and TMS (TMS vs. NoTMS) were performed for each ROI (M1, PMC, and SMA) according to task conditions. All results were summarised in Fig. 2B. During LHC, there was a significant main effect of hemisphere in all ROIs (M1: F1, 11 = 226.98, p < 0.001; PMC: F1, 11 = 160.22, p < 0.001; SMA: F1, 11 = 53.58, p < 0.001) and a significant TMS effect in the SMA (F1, 11 = 6.50, p < 0.05). The M1 (F1,11 = 3.82, p = 0.08) and the PMC (F1, 11 = 3.84, p = 0.08) showed a marginally significant TMS effect. There was a significant interaction between hemisphere × TMS in the M1 (F1, 11 = 19.96, p < 0.001) and between hemisphere × site × TMS in the M1 (F1, 11 = 10.79, p < 0.01) and the PMC (F1, 11 = 15.10, p < 0.01). *Post-hoc* comparisons between TMS and NoTMS phases showed that the left M1 TMS evoked significant reduction in the regional activity in the right (contralateral) hemisphere, whereas the right M1 TMS induced a significant increase of the magnitude of deactivation in the left motor areas. There was no TMS effect in the ROIs at the stimulated (ipsilateral) hemisphere. During RHC, the motor regions showed the similar pattern of activity to the LHC condition. There was a significant main effects of hemisphere (M1: F1, 11 = 176.48, p < 0.001; PMC: F1, 11 = 81.43, p < 0.001; SMA: F1, 11 = 48.80, p < 0.001) and TMS (M1: F1, 11 = 5.64, p < 0.05; PMC: F1, 11 = 15.11, p < 0.01; SMA: F1, 11 = 11.11, p < 0.01). The 3 way interaction of hemisphere × site × TMS was significant for all ROIs (M1: F1, 11 = 14.06, p < 0.01; PMC: F1, 11 = 23.50, p < 0.001; SMA: F1, 11 = 8.58, p < 0.05). *Post-hoc* comparisons between TMS and NoTMS phase revealed the similar findings to the results of the LHC: the left M1 TMS evoked the significant increase of deactivations in the contralateral (right) hemisphere, whereas right M1 TMS induced significant decreases of activity in all contralateral ROIs. During BHC, the ANOVAs revealed a significant main effect of TMS (M1: F1, 11 = 11.51, p < 0.01; PMC: F1, 11 = 6.67, p < 0.05; SMA: F1, 11 = 10.48, p < 0.01). Only the M1 showed a significant main effect of hemisphere (M1: F1, 11 = 7.92, p < 0.001). There was a significant interaction between hemisphere × site × TMS in the M1 (F1, 11 = 12.77, p < 0.01) and SMA (F1, 11 = 4.75, p = 0.05). *Post-hoc* comparisons between TMS and NoTMS phases showed that the left M1 TMS evoked a significant reduction in the activity of the contralateral (right) M1 and SMA, whereas the right M1 TMS induced a significant decrease in the activity of the left M1, PMC, as well as the right SMA. In the MN, we demonstrated that a short train of 1Hz TMS to the M1 induced a strong interhemispheric inhibition resulting in decreased activation/increased deactivation in the contralateral cortical regions across the task conditions.

To investigate the M1 TMS effect in the whole brain, we performed a conjunction analysis by comparing TMS with NoTMS phase across the task conditions. Fig. 2C summarises the results. M1 TMS evoked significant activation in the bilateral secondary somatosensory cortex (SII) including operculum and superior temporal gyrus (STG) as well as deactivation in the contralateral precentral gyrus/postcentral gyrus (M1/S1), middle frontal gyrus (MFG) and bilateral SPL, precuneus, and middle occipital gyrus (MOG). A repeated-measures ANOVA with task (LHC, RHC, BHC and rest) and TMS site (left vs. right) was performed in these regions. The ANOVAs revealed that a significant effect of task for the all ROIs (SII [operculum]: F3, 9 = 33.13, p < 0.001; SII [STG]: F3, 9 = 26.85, p < 0.001; precuneus: F3, 9 = 13.60, p < 0.001; SPL: F3, 9 = 11.31, p < 0.001). The other main and interaction effects were not significant (ps > 0.2). Subsequent t-tests for the overlapping regions demonstrated that the TMS effects were stronger at rest than any other task conditions.

Without the motor execution task, the left M1 stimulation evoked the significant activation in the bilateral STG/SII and deactivation in superior occipital gyrus (SOG) and precuneus (Fig. 2D & Table S2). The vertex stimulation caused significant deactivation in the medial prefrontal cortex (mPFC), superior frontal gyrus (SFG), middle frontal gyrus (MFG), precuneus, and visual cortex (Fig. 2E & Table S2).

### ICA results

ICA revealed 13 components showing the patterns of temporally coherent signal in the brain. Fig. 3 and Table S3 summarises 13 functional brain networks. Fig. 4 displays the result of the regression analysis on each network, showing how the different networks were modulated by the experimental paradigms in the 3 groups.

**Fig. 3.**
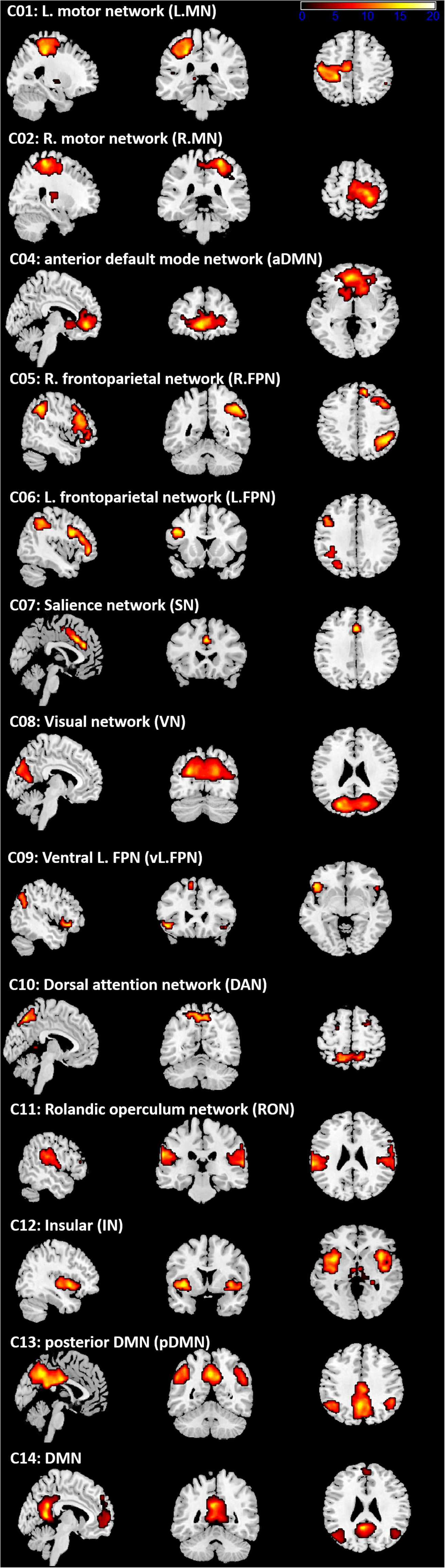
ICA results. Spatial distribution of 13 networks. See Table 2 for coordinates.

**Fig. 4.**
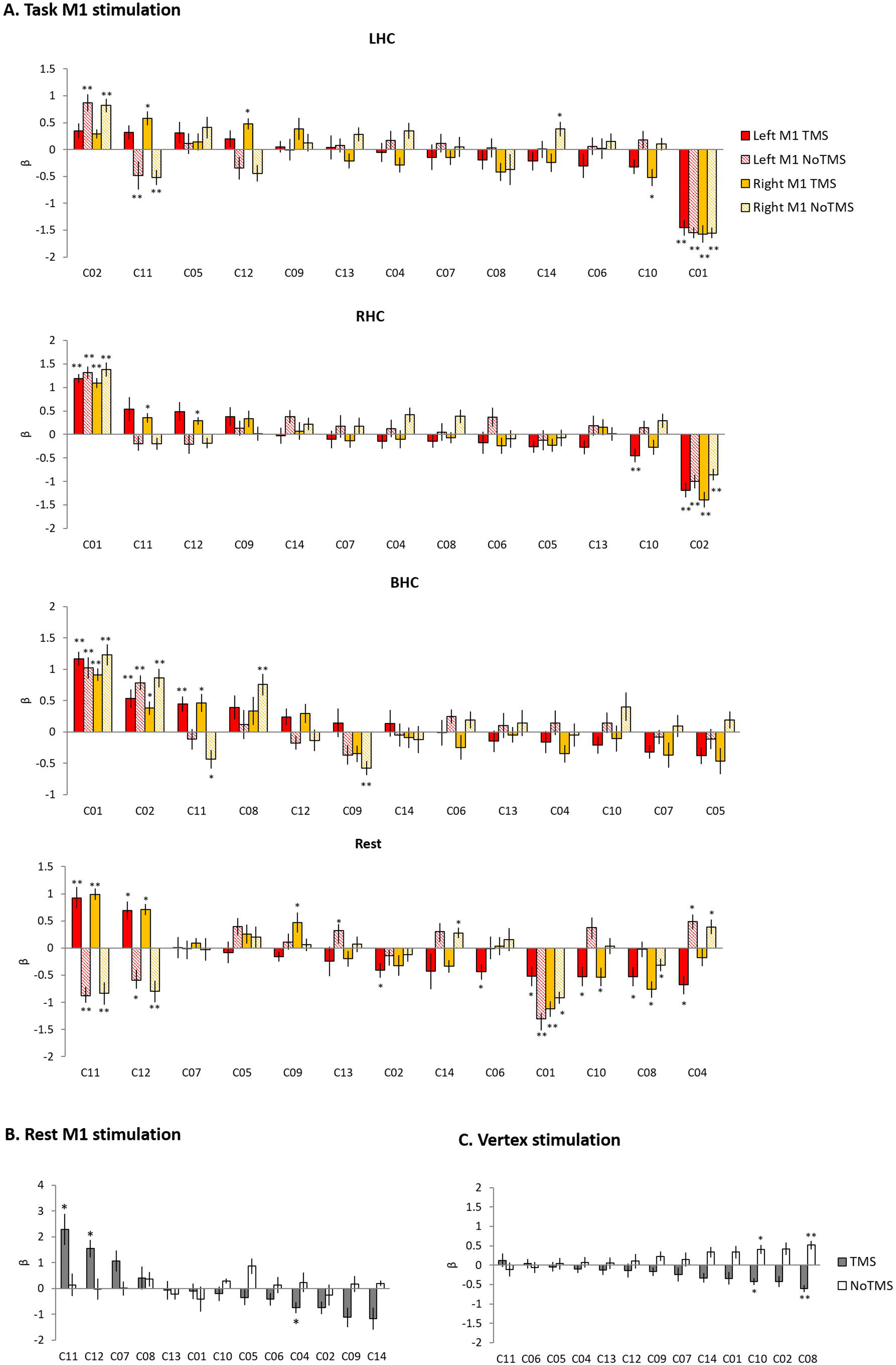
The results of temporal regression analysis. Bar chart showing the mean β value for each condition from a regression analysis performed on each of the 13 networks. A) Task M1 stimulation. B) Rest M1 stimulation. C) Vertex stimulation. Error bars indicate standard errors. ** p _FDR-corrected_ < 0.005, * p _FDR-corrected_ < 0.05

In the task M1 stimulation group (Fig. 3A), we found several networks were significantly modulated by task and TMS: C01 (L.MN). C02 (R.MN), C04 (aDMN), C08 (VN), C09 (vl.FPN), C10 (DAN), C11 (RON), and C12 (IN). C01 consisted with the left M1, S1, SMA, RO, insular, putamen and thalamus and C02 included the right M1, S1, SMA, and RO. C04 was composed of the mPFC, anterior cingulate cortex (ACC) and superior medial gyrus (SMG). C08 consisted of the primary visual cortex. C09 included the left IFG, angular gyrus (AG) and inferior parietal lobe (IPL). C10 consisted of the bilateral frontal eye field and SPL. C11 contained the bilateral RO and STG and C12 was composed of the bilateral insular. The LHC significantly activated C02 (R.MN) without TMS, C11 (RON) and C12 (IN) with the right M1 TMS, whereas deactivated C01 (L.MN), C11 without TMS and C10 (DAN) with the right M1 TMS. The RHC showed the significant activation in C01 (L.MN) across the all conditions, C11 (RON) and C12 (IN) with the right M1 TMS as well as the deactivation in C02 (R.MN) and C10 (DAN) with the left M1 TMS. The BHC significantly activated C01 (L.MN) and C02 (R.MN) regardless of the conditions. During BHC, C11 (RON) showed the significant activation with TMS and deactivation during the right M1 without TMS. C09 (vl.FPN) was deactivated with the right M1 without TMS. The rest condition showed different pattern of network modulation compared to motor execution condition. TMS significantly activated C11 (RON) and C12 (IN), whereas deactivated C08 (VN) and C10 (DAN) regardless of the site as well as C02 (R.MN), C04 (aDMN), C06 (L.FPN) with the left M1 TMS. C09 (vl.FPN) showed the significant activation with the right M1 TMS.

The rest M1 stimulation group showed that TMS significantly activated C11 and C12 and deactivated C04 and C14 (DMN) (Fig. 4B). C14 consisted of the mPFC, precuneus and bilateral AG. The vertex group showed the deactivation in C10 and C08 (Fig. 4C).

To investigate the effect of the M1 stimulation at work and rest, we selected the networks significantly modulated by task and TMS (C01, C02, C04, C08, C09, C10, C11, C12 and C14). First, for the task M1 stimulation group, we used 2 × 2 ANOVA with task (LHC, RHC, BHC and rest) and TMS (TMS and noTMS) as within-subject factors to examine the effect of M1 stimulation during motor execution task. Fig. 5A displays the results of task-specific networks (C01 [L.MN] and C02 [R.MN]). C01 showed a significant main effect of task (F_1,11_ = 62.48, p < 0.001) and an interaction (F_3,9_ = 4.70, p = 0.031). *Post-hoc* t-tests revealed that the left M1 TMS significantly reduced the deactivation at rest. C02 showed the main effect of task (F_1,11_ = 28.21, p < 0.001) and TMS (F_1,11_ = 4.99, p = 0.047). *Post-hoc* t-tests revealed that the left M1 TMS reduced the activity during LHC. The right M1 TMS also decreased the activation during LHC and BHC, whereas increased deactivation during RHC. Fig. 5B shows the result of the M1 TMS-specific networks (C11 [RON] and C12 [IN]). C11 showed a significant main effect of TMS (F_1,11_ = 32.11, p < 0.001) and an interaction (F_3,9_ = 6.13, p = 0.015). *Post-hoc* t-tests revealed that the M1 TMS significantly increased the network activity across all task conditions. C12 showed the significant main effect of TMS (F_1,11_ = 24.87, p < 0.001). *Post-hoc* t-tests revealed that the M1 TMS significantly increased the network activity across all task conditions.

**Fig. 5.**
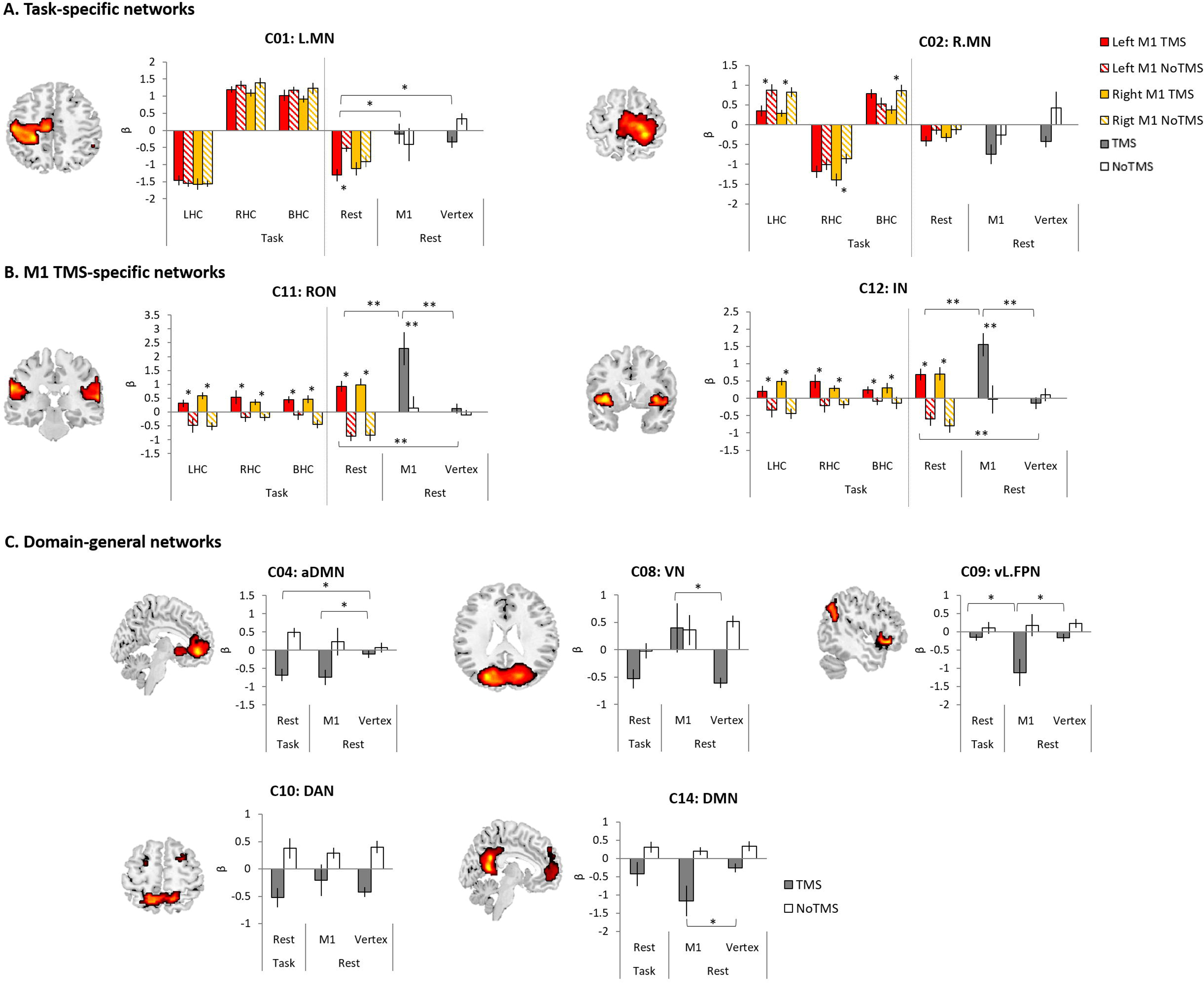
The results of temporal regression analysis between groups. A) Task-specific networks. B) M1 TMS specific networks. C) Domain-general networks. The bar filled colour indicates the TMS phase and the bar filled with diagonal lines indicates the No TMS phase. Red (left M1 TMS) and Yellow (right M1 TMS). Grey bar indicated TMS phase and white bar indicate NoTMS phase. Error bars indicate standard errors. ** p < 0.005, * p < 0.05

Second, to compare the effect of the M1 stimulation during the rest condition within the motor execution task and at rest, ANOVA with TMS (TMS and noTMS) as a within-subject factor and group (task M1, rest M1 and vertex) as a between-subject factor was conducted. The rest condition during motor task was the left M1 TMS session. C01 showed the main effect of group (F_2,33_ = 23.61, p < 0.001), whereas C02 showed the main effect of TMS (F_1,33_ = 5.87, p = 0.021) and group (F_2,33_ = 4.83, p = 0.014). *Post-hoc* t-tests on the TMS phase between groups revealed that the task M1 stimulation group showed the significant deactivation in C01 compared to the rest M1 and vertex (Fig. 5A). The M1 TMS-specific networks showed the significant main effect of TMS (C11: F_1,33_ = 31.22, p < 0.001; C12: F_1,33_ = 15.92, p < 0.001) and group (C11: F_2,33_ = 7.51, p = 0.002; C12: F_2,33_ = 5.62, p = 0.008) as well as the interaction (C11: F_2,33_ = 5.50, p = 0.009; C12: F_2,33_ = 6.67, p = 0.004). *Post-hoc* t-tests on the TMS phase demonstrated that the rest M1 stimulation evoked the stronger networks activation in both C11 and C12 compared to other groups and the task M1 stimulation also showed stronger activation in both networks than vertex group (Fig. 5B). Other domain-general networks were examined to detect the effect of M1 stimulation (Fig. 5C). C04 showed a significant main effect of TMS (F_1,33_ = 19.68, p < 0.001) and a marginally significant interaction (F_2,33_ = 3.09, p = 0.059). *Post-hoc* t-tests on the TMS phase revealed that both task M1 and rest M1 stimulation significantly decreased the network deactivation relative to the vertex stimulation. C08 showed a main effect of TMS (F_1,33_ = 6.81, p = 0.014) and group (F_2,33_ = 4.03, p = 0.027). *Post-hoc* t-tests on the TMS phase revealed that the rest M1 stimulation evoked the increased activation more than the vertex stimulation. C09 showed a main effect of TMS (F_1,33_ = 14.86, p = 0.001) and an interaction (F_2,33_ = 3.69, p = 0.036). *Post-hoc* t-tests on the TMS phase revealed that rest M1 stimulation significantly deactivated the network activity compared to other groups. We found the main effect of TMS in C10 (F_1,33_ = 22.41, p < 0.001). C14 showed a main effect of TMS (F_1,33_ = 19.68, p = 0.001) a marginally significant interaction (F_2,33_ = 2.97, p = 0.069). *Post-hoc* t-tests on the TMS phase revealed that the rest M1 stimulation significantly deactivated the network compared to the vertex stimulation.

To detect the FNC changes modulated by TMS and task, we conducted a One-way ANOVA with group (task M1 [left M1 TMS], rest M1 and vertex) on the FNC between the networks showed the effect of them in the previous analysis. We found a significant group effect in the FNC between these networks (Fig 6A and Table S4). *Post hoc t-tests* between groups demonstrated that the task M1 group compared to the rest M1 group showed the significantly decreased FNC in C01-C02, C09-C10 and C10-C14 as well as the increased FNC in C04-C09 (Fig 6B left). The rest M1 stimulation relative to the vertex stimulation showed the decreased FNC in C02-C14, C04-C12, C08-C12, C08-C14 and C11-C14 (Fig 6B right).

**Fig. 6.**
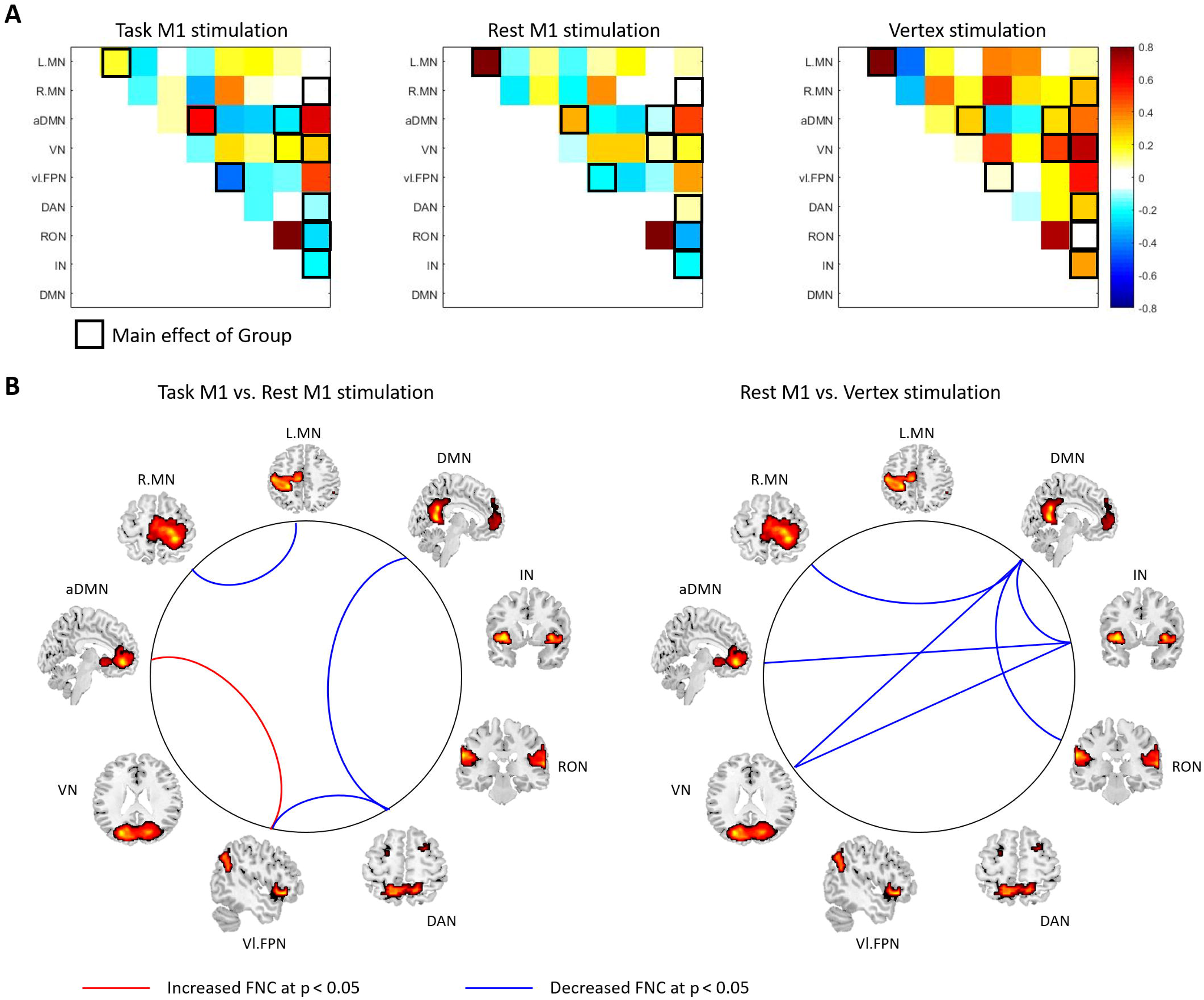
The results of FNC analysis between groups. A) The FNC matrix. Pairwise correlation coefficients between RSN time courses were Fisher z-transformed and averaged across subjects within the task M1 [left M1 TMS session], rest M1 and vertex stimulation groups. The black boxes indicate the FNC that showed a significant group effect (p < 0.05). Warm colours indicates the positive coupling and cool colours indicates the negative coupling (decoupling) between the networks. B) The FNC comparisons between two groups (Left: Task M1 vs. Rest M1 stimulation; Right: Rest M1 vs. Vertex stimulation). Red line indicates the increased FNC and blue line shows the decreased FNC between the groups (p < 0.05).

## Discussion

A novel finding of this study is that there were two non-motor networks specifically modulated by the M1 stimulation: RON (C11) and IN (C12). The M1 stimulation evoked activation in these networks during task and at rest. The RO is a part of the SII (Eickhoff et al., 2006) and involved in a wide range of somatic stimuli processing (Ledberg et al., 1995;Roland et al., 1998;Bodegard et al., 2000). Anatomically, this area is directly connected to key multisensory regions including PMC, M1, IPL, inferior parietal sulcus and inferior frontal cortices and functionally associated with the multisensory perception of the hand through the collaboration with the frontoparietal regions (Eickhoff et al., 2010;Gentile et al., 2011). TMS over the M1 induces peripheral muscle activity at the targeted hand. A previous study with supratheshold TMS during fMRI have shown that this afferent feedback caused by TMS can contribute to BOLD signal changes in M1 and somatosensory areas (Denslow et al., 2005). However, the M1 TMS generates not only physical muscle responses but also the awareness of unintended muscle activity. Participants experiences that their hand was strained or sometimes twitched by the M1 stimulation, not by themselves, which alerts the sense of ownership or agency for one’s hand and its movements (Murray and Wallace, 2012). Thus, this unique experience of the M1 stimulation may drive bodily self-consciousness (BSC) – the processing and integration of multisensory bodily signals (Lenggenhager et al., 2007;Blanke, 2012). The RO has been repeatedly associated with BSC, showing increased activation in interoceptive signals from the body parts (Damasio and Meyer, 2009;Blefari et al., 2017). The strong regional activity and the increased network activation in the RO may be attributed to not only the afferent feedback from stimulating the M1 but also the BSC.

In addition to the RO, the insular play a critical role in BSC (Damasio and Meyer, 2009;Blanke, 2012;Gogolla, 2017). The insular is located deep within the lateral sulcus of each hemisphere, heavily connected to cortical and subcortical regions serving sensory, emotion and cognitive functions (Shelley and Trimble, 2004). The insular receives information from outside the body (auditory, somatosensory, olfactory, gustatory and visual information) and from inside the body (interoceptive information). Based on anatomical and functional connectivity of the insular, Craig suggested that the insula is a locus to form the self-awareness of feelings from the body (Craig, 2011). The M1 stimulation induced strong activation in the IN during task and at rest compared to the vertex stimulation. Similar to the RO, the involvement of the IN can be attributed to the self-awareness of feeling caused by the M1 TMS. Furthermore, the RON and IN showed the decoupling with the DMN when the TMS was applied over the M1 compared to the vertex. The DMN is a network shows task-related deactivation and involved in internally focused processing such as mind wandering and consciousness (Buckner et al., 2008). It suggests that the unique multisensory experience of the M1 stimulation prompted the involvement of these networks by disrupting on-going internal processing.

We found that the pattern of network activation in the RON and IN was state-dependent: stronger activation at rest and even in the rest condition during motor execution. Previously, Bestmann et al (2008) demonstrated state-dependent TMS effect in the motor system. The finding indicated that the effects of the TMS is highly dependent on the existing state of activity of the stimulated area. Here, we showed the similar finding in non-targeted neural networks. It might be possible that the goal-oriented task processing disturbed the interoceptive processing of the M1 TMS experience, leading to the reduced the network activation during task.

In the motor system, we found that a short train of 1Hz TMS to the M1 induced a strong interhemispheric inhibition during motor execution at the regional activity as well as the FNC between the left and right motor networks. The interhemispheric inhibition caused by TMS has been widely investigated in both humans and animals (Chang, 1953;Matsunami and Hamada, 1984;Ferbert et al., 1992;Wassermann et al., 1998;Hanajima et al., 2001) suggesting that the interhemispheric inhibition is mediated through transcallosal pathways (Meyer et al., 1995;1998). Recently, studies combining TMS with neuroimaging (PET/fMRI) have showed that TMS over the M1 modulated brain activity in the motor areas connected to the targeted M1 (Bohning et al., 1999;2000a;2000b;Bestmann et al., 2003;2004). In accordance with previous studies, our ROI results demonstrated that the M1 stimulation induced the interhemispheric inhibition, decreasing the task-related activation and increasing the deactivation at the contralateral motor regions. However, at the network level, we found that the TMS effect in the MN of the dominant hemisphere (left) was not evident as that in the non-dominant MN (R.MN). R.MN showed the significant TMS effect, showing the reduction of network activation and the enlargement of network deactivation. Contrary to R.MN, M1 TMS did not modulate L.MN during the task but the left M1 TMS at rest increased the network deactivation. It might be possible that the current TMS paradigm (11 pulses with 100% RMT) was not enough to elicit neural changes at the network level when the network linked to the dominant hand. It is noted that our participants were strong right-handed. This finding is compatible with previous reports demonstrating the absence of significant alterations in brain activity at the M1 after TMS (Bohning et al., 2000a;Baudewig et al., 2001). Bohning et al (2000a) tested the effects of TMS on M1 during a simple motor execution task. They delivered 1 Hz TMS with 110% RMT over the M1 and estimated the level of regional activity at the target region. The regional activity associated with the task and TMS was not different from the activity evoked by the task alone.

We showed that the M1 stimulation deactivated other domain-general networks. Previously, we demonstrated that the M1 stimulation inhibited the DMN compared to the vertex stimulation (Jung et al., 2016). Similarly, we found that the M1 stimulation deactivated aDMN (C04) and DMN (C14) relative to the vertex stimulation. The experience of the M1 TMS disrupts the internally focused processing resulting the deactivation of the DMN (Jung et al., 2016). A task-active network (C09) showed state-dependent TMS effect. The vl.FPN (C09) is a sub-set of the FPN contributing to executive processing across tasks (Fedorenko et al., 2013). Although the M1 TMS deactivated the network at rest not during task, the network was not involved in any task conditions during motor execution. It might be driven from uncontrolled mental activity at rest, not directly connected to M1 stimulation. The DAN (C10) showed the deactivation when TMS was delivered to the M1 or vertex. The DAN is involved in the top-down guided voluntary allocation of attention to sensory input (Vossel et al., 2014). The general by-product of TMS pulse such as tactile sensation and ‘click’ sound potentially contributed to the deactivation of the network. In addition to the TMS effect, task also modulated the interaction between these networks – decoupling task-active network (DAN) and task-negative network (DMN).

To our best knowledge, this study is the first study to examine multiple brain networks associated with the unique experience of the M1 stimulation. We demonstrated that the M1 stimulation modulated the targeted motor system as well as other non-motor, remote functional brain networks. Especially, functional networks related to BSC showed the M1 TMS specific modulation responding to its multisensory experience. Our results support that the state of brain is a critical factor to influence the effect of the M1 stimulation in these networks. Our findings shed light on understanding the neural mechanism of the complex and multisensory experience of the M1 stimulation.

## Author Contribution

J.J and S.J designed the study. A.B and R.R developed the concurrent TMS/fMRI system. J.J and A.B collected the data. J.J analysed the data. J.J and S.J wrote the manuscript and approved the final manuscript.

## Funding

This work was supported by the University of Nottingham’s research fellowship programme to J.J, the National Research Foundation of Korea’s WCU (World Class University) programme funded by the Ministry of Education, Science and Technology (R31-2008-000-10008-0) and by an MRC programme grant G0901321 awarded to RB and SRJ.

## Conflict of Interest

None.

## References

Baudewig, J., Siebner, H.R., Bestmann, S., Tergau, F., Tings, T., Paulus, W., and Frahm, J. (2001). Functional MRI of cortical activations induced by transcranial magnetic stimulation (TMS). Neuroreport 12, 3543–3548.

Bell, A.J., and Sejnowski, T.J. (1995). An information-maximization approach to blind separation and blind deconvolution. Neural Comput 7, 1129–1159.

Bestmann, S., Baudewig, J., Siebner, H.R., Rothwell, J.C., and Frahm, J. (2003). Subthreshold high-frequency TMS of human primary motor cortex modulates interconnected frontal motor areas as detected by interleaved fMRI-TMS. Neuroimage 20, 1685–1696.

Bestmann, S., Baudewig, J., Siebner, H.R., Rothwell, J.C., and Frahm, J. (2004). Functional MRI of the immediate impact of transcranial magnetic stimulation on cortical and subcortical motor circuits. Eur J Neurosci 19, 1950–1962.

Bestmann, S., Baudewig, J., Siebner, H.R., Rothwell, J.C., and Frahm, J. (2005). BOLD MRI responses to repetitive TMS over human dorsal premotor cortex. Neuroimage 28, 22–29.

Bestmann, S., Swayne, O., Blankenburg, F., Ruff, C.C., Haggard, P., Weiskopf, N., Josephs, O., Driver, J., Rothwell, J.C., and Ward, N.S. (2008). Dorsal premotor cortex exerts state-dependent causal influences on activity in contralateral primary motor and dorsal premotor cortex. Cereb Cortex 18, 1281–1291.

Blanke, O. (2012). Multisensory brain mechanisms of bodily self-consciousness. Nat Rev Neurosci 13, 556–571.

Blefari, M.L., Martuzzi, R., Salomon, R., Bello-Ruiz, J., Herbelin, B., Serino, A., and Blanke, O. (2017). Bilateral Rolandic operculum processing underlying heartbeat awareness reflects changes in bodily self-consciousness. Eur J Neurosci 45, 1300–1312.

Bodegard, A., Ledberg, A., Geyer, S., Naito, E., Zilles, K., and Roland, P.E. (2000). Object shape differences reflected by somatosensory cortical activation. J Neurosci 20, RC51.

Bohning, D., Shastri, A., Lomarev, M., Nahas, Z., and George, M. (2003). BOLD-fMRI vs. transcranial magnetic stimulation (TMS) pulse-train length: testing for linearity. J Magn Reson Imaging 17, 279–290.

Bohning, D., Shastri, A., Mcconnell, K., Nahas, Z., Lorberbaum, J., Robert, D., Teneback, C., Vincent, D., and George, M. (1999). A combined TMS/fMRI study of intensity-dependent TMS over motor cortex. Biol Psychiatry 45, 385–394.

Bohning, D., Shastri, A., Mcgavin, L., Mcconnell, K., Nahas, Z., Lorberbaum, J., Robert, D., and George, M. (2000a). Motor cortex brain activity induced by 1-Hz transcanial magnetic stimulation is stimilar in location and level to that for volitional movement. Invest Radiol 35, 676–683.

Bohning, D., Shastri, A., Nahas, Z., Lorberbaum, J., Andersen, S., Danels, W., Haxthausen, E., Vincent, D., and George, M. (1998). Echo-planar BOLD fMRI of brain acitvation induced by concurrent transcanial magnetic stimulation. Invest Radiol 33, 336–340.

Bohning, D., Shastri, A., Wassermann, E., Ziemann, U., Lorberbaum, J., Nahas, Z., Lomarev, M., and George, M. (2000b). BOLD-fMRI response to single-pulse transcanial magnetic stimulation (TMS). J Magn Reson Imaging 11, 569–574.

Buckner, R.L., Andrews-Hanna, J.R., and Schacter, D.L. (2008). The brain’s default network: anatomy, function, and relevance to disease. Ann N Y Acad Sci 1124, 1–38.

Calhoun, V.D., Adali, T., Pearlson, G.D., and Pekar, J.J. (2001). A method for making group inferences from functional MRI data using independent component analysis. Hum Brain Mapp 14, 140–151.

Chang, H.T. (1953). Cortical response to activity of callosal neurons. J Neurophysiol 16, 117–131.

Craig, A.D. (2011). Significance of the insula for the evolution of human awareness of feelings from the body. Ann N Y Acad Sci 1225, 72–82.

Damasio, A., and Meyer, K. (2009). Consciousness: An Overview of the Phenomenon and of Its Possible Neural Basis. Neurology of Consciousness: Cognitive Neuroscience and Neuropathology, 3–14.

Denslow, S., Lomarev, M., George, M.S., and Bohning, D.E. (2005). Cortical and subcortical brain effects of transcranial magnetic stimulation (TMS)-induced movement: an interleaved TMS/functional magnetic resonance imaging study. Biol Psychiatry 57, 752–760.

Di Lazzaro, V., Oliviero, A., Profice, P., Insola, A., Mazzone, P., and Tonali, P. (1999). Direct demonstration of interhemispheric inhibition of the human motor cortex produced by transcranial magnetic stimulation.. Experimental Brain Research 124, 520–524.

Egolf, E.A., Calhoun, V.D., and Kiehl, K.A. (2004). Group ICA of fMRI Toolbox (GIFT). Biological Psychiatry 55, 8S–8S.

Eickhoff, S.B., Jbabdi, S., Caspers, S., Laird, A.R., Fox, P.T., Zilles, K., and Behrens, T.E. (2010). Anatomical and functional connectivity of cytoarchitectonic areas within the human parietal operculum. J Neurosci 30, 6409–6421.

Eickhoff, S.B., Schleicher, A., Zilles, K., and Amunts, K. (2006). The human parietal operculum. I. Cytoarchitectonic mapping of subdivisions. Cereb Cortex 16, 254–267.

Fedorenko, E., Duncan, J., and Kanwisher, N. (2013). Broad domain generality in focal regions of frontal and parietal cortex. Proc Natl Acad Sci U S A 110, 16616–16621.

Ferbert, A., Priori, A., Rothwell, J.C., Day, B.L., Colebatch, J.G., and Marsden, C.D. (1992). Interhemispheric inhibition of the human motor cortex. J Physiol 453, 525–546.

Gentile, G., Petkova, V.I., and Ehrsson, H.H. (2011). Integration of visual and tactile signals from the hand in the human brain: an FMRI study. J Neurophysiol 105, 910–922.

Gogolla, N. (2017). The insular cortex. Curr Biol 27, R580–R586.

Hanajima, R., Ugawa, Y., Machii, K., Mochizuki, H., Terao, Y., Enomoto, H., Furubayashi, T., Shiio, Y., Uesugi, H., and Kanazawa, I. (2001). Interhemispheric facilitation of the hand motor area in humans. J Physiol 531, 849–859.

Himberg, J., Hyvarinen, A., and Esposito, F. (2004). Validating the independent components of neuroimaging time series via clustering and visualization. Neuroimage 22, 1214–1222.

Jafri, M.J., Pearlson, G.D., Stevens, M., and Calhoun, V.D. (2008). A method for functional network connectivity among spatially independent resting-state components in schizophrenia. Neuroimage 39, 1666–1681.

Jung, J., Bungert, A., Bowtell, R., and Jackson, S.R. (2016). Vertex Stimulation as a Control Site for Transcranial Magnetic Stimulation: A Concurrent TMS/fMRI Study. Brain Stimul 9, 58–64.

Koch, G., Franca, M., Albrecht, U.-V., Caltagirone, C., and Rothwell, J. (2006). Effects of paired pulse tms of primary somatosensory cortex on perception of a peripheral electrical stimulus. Experimental Brain Research 172, 416–424.

Ledberg, A., O’sullivan, B.T., Kinomura, S., and Roland, P.E. (1995). Somatosensory activations of the parietal operculum of man. A PET study. Eur J Neurosci 7, 1934–1941.

Lenggenhager, B., Tadi, T., Metzinger, T., and Blanke, O. (2007). Video ergo sum: manipulating bodily self-consciousness. Science 317, 1096–1099.

Li, X., Teneback, C.C., Nahas, Z., Kozel, F.A., Large, C., Cohn, J., Bohning, D.E., and George, M.S. (2004). Interleaved transcranial magnetic stimulation/functional MRI confirms that lamotrigine inhibits cortical excitability in healthy young men. Neuropsychopharmacology 29, 1395–1407.

Matsunami, K., and Hamada, I. (1984). Effects of stimulation of corpus callosum on precentral neuron activity in the awake monkey. J Neurophysiol 52, 676–691.

Meyer, B.U., Roricht, S., Grafin Von Einsiedel, H., Kruggel, F., and Weindl, A. (1995). Inhibitory and excitatory interhemispheric transfers between motor cortical areas in normal humans and patients with abnormalities of the corpus callosum. Brain 118 (Pt 2), 429–440.

Meyer, B.U., Roricht, S., and Woiciechowsky, C. (1998). Topography of fibers in the human corpus callosum mediating interhemispheric inhibition between the motor cortices. Ann Neurol 43, 360–369.

Murray, M.M., and Wallace, M.T. (2012). The neural bases of multisensory processes. Boca Raton: CRC Press.

Oldfield, R.C. (1971). The assessment and analysis of handedness: the Edinburgh inventory. Neuropsychologia 9, 97–113.

Roland, P.E., O’sullivan, B., and Kawashima, R. (1998). Shape and roughness activate different somatosensory areas in the human brain. Proc Natl Acad Sci U S A 95, 3295–3300.

Ruff, C.C., Blankenburg, F., Bjoertomt, O., Bestmann, S., Weiskopf, N., and Driver, J. (2009). Hemispheric differences in frontal and parietal influences on human occipital cortex: direct confirmation with concurrent TMS-fMRI. J Cogn Neurosci 21, 1146–1161.

Shastri, A., George, M.S., and Bohning, D.E. (1999). Performance of a system for interleaving transcranial magnetic stimulation with steady-state magnetic resonance imaging. Electroencephalogr Clin Neurophysiol Suppl 51, 55–64.

Shelley, B.P., and Trimble, M.R. (2004). The insular lobe of Reil--its anatamico-functional, behavioural and neuropsychiatric attributes in humans--a review. World J Biol Psychiatry 5, 176–200.

Steinmetz, H., Furst, G., and Meyer, B.U. (1989). Craniocerebral topography within the international 10–20 system. Electroencephalogr Clin Neurophysiol 72, 499–506.

Vossel, S., Geng, J.J., and Fink, G.R. (2014). Dorsal and ventral attention systems: distinct neural circuits but collaborative roles. Neuroscientist 20, 150–159.

Walsh, V., and Cowey, A. (2000). Transcaniall magnetic stimulation and cognitive neuroscience. Nature Reviews Neuroscience 1, 73–80.

Wasserman, E., Epstein, C.M., and Ziemann, U. (2008). The Oxford handbook of transcranial stimulation. Oxford ; New York: Oxford University Press.

Wassermann, E.M., Wedegaertner, F.R., Ziemann, U., George, M.S., and Chen, R. (1998). Crossed reduction of human motor cortex excitability by 1-Hz transcranial magnetic stimulation. Neurosci Lett 250, 141–144.

