## Supplemental information for "Modulating brain networks with transcranial magnetic stimulation over the primary motor cortex: a concurrent TMS/fMRI study"

Supplementary Table 1

| Task M1 stimulation | | | | | | |
| --- | --- | --- | --- | --- | --- | --- |
|  | x | y | z | x | y | z |
| Sub01 | -40 | -12 | 66 | 34 | -8 | 68 |
| Sub02 | -21 | -30 | 74 | 16 | -14 | 78 |
| Sub03 | -34 | -18 | 70 | 34 | -4 | 68 |
| Sub04 | -36 | -24 | 70 | 36 | -2 | 66 |
| Sub05 | -36 | -4 | 66 | 35 | -10 | 60 |
| Sub06 | 40 | -26 | 68 | 30 | 0 | 70 |
| Sub07 | -30 | -42 | 74 | 38 | -18 | 68 |
| Sub08 | -38 | -16 | 68 | 22 | -12 | 74 |
| Sub09 | -40 | -22 | 68 | 36 | -10 | 68 |
| Sub10 | -36 | -18 | 70 | 36 | -14 | 68 |
| Sub11 | -30 | -24 | 72 | 32 | -24 | 74 |
| Sub12 | -30 | -2 | 70 | 30 | -4 | 68 |
| Rest M1 stimulation | | | | | | |
| Sub01 | -34 | -26 | 72 |  |  |  |
| Sub02 | -52 | -26 | 60 |  |  |  |
| Sub03 | -26 | -32 | 76 |  |  |  |
| Sub04 | -34 | -4 | 68 |  |  |  |
| Sub05 | -38 | 16 | 60 |  |  |  |
| Sub06 | -24 | -54 | 74 |  |  |  |
| Sub07 | -36 | -26 | 72 |  |  |  |
| Sub08 | -30 | -38 | 72 |  |  |  |
| Sub09 | -28 | -30 | 74 |  |  |  |
| Sub10 | -36 | -4 | 66 |  |  |  |
| Sub11 | -36 | -24 | 70 |  |  |  |
| Sub12 | -38 | -16 | 68 |  |  |  |
| Vertex stimulation | | | | | | |
| Sub01 | 6 | -12 | 76 |  |  |  |
| Sub02 | 0 | -44 | 72 |  |  |  |
| Sub03 | -4 | -52 | 72 |  |  |  |
| Sub04 | 2 | -10 | 70 |  |  |  |
| Sub05 | 3 | -15 | 72 |  |  |  |
| Sub06 | -2 | -20 | 73 |  |  |  |
| Sub07 | 5 | -16 | 74 |  |  |  |
| Sub08 | -5 | -13 | 75 |  |  |  |
| Sub09 | 0 | -15 | 70 |  |  |  |
| Sub10 | -3 | 5 | 75 |  |  |  |
| Sub11 | 1 | 2 | 76 |  |  |  |
| Sub12 | -1 | 1 | 73 |  |  |  |

Table S1. The MNI coordinates of TMS target site

Supplementary Table 2

| Group | Contrasts | Cluster region | Cluster extent | Peak MNI coordinate | | |
| --- | --- | --- | --- | --- | --- | --- |
|  |  |  |  | x | y | z |
| Task M1 stimulation | Main effect of task | M1 | 1273 | -33 | -24 | 63 |
|  |  | S1 |  | -36 | -24 | 51 |
|  |  | PMC |  | -50 | -15 | 37 |
|  |  | SMA |  | -3 | -6 | 48 |
|  |  | M1 | 770 | 36 | -15 | 54 |
|  |  | PMC |  | 52 | -12 | 38 |
|  |  | SMA |  | 6 | -6 | 48 |
|  |  | Thalamus | 450 | -15 | -27 | 0 |
|  |  | Putamen |  | -30 | -15 | -3 |
|  |  | RO |  | -39 | -21 | 18 |
|  |  | Thalamus | 356 | 15 | -24 | 0 |
|  |  | Pallidum |  | 30 | -9 | -6 |
|  |  | RO |  | 39 | -18 | 15 |
|  |  | MOG | 88 | -18 | -90 | 15 |
|  |  | SOG |  | -15 | -81 | 30 |
|  |  | MCC | 69 | 12 | -6 | 45 |
|  |  | Calcarine gyrus | 69 | 15 | -84 | 9 |
|  | Main effect of site | Calcarine gyrus | 354 | 18 | -51 | 12 |
|  |  | STG |  | 51 | -45 | 12 |
|  |  | MTG |  | 66 | -45 | 6 |
|  |  | Lingual gyrus |  | 18 | -54 | 3 |
|  |  | S1 | 325 | -36 | -30 | 54 |
|  |  | IPL |  | -45 | -27 | 45 |
|  |  | SPL |  | -33 | -45 | 57 |
|  |  | IFG | 324 | 51 | 30 | -6 |
|  |  | Putamen |  | 27 | 15 | 9 |
|  |  | M1 | 202 | 36 | -12 | 51 |
|  |  | S1 | 179 | 42 | -30 | 57 |
|  |  | M1 | 125 | -36 | -9 | 48 |
|  |  | Calcarine gyrus | 109 | -15 | -78 | 15 |
|  |  | Cuneus |  | 3 | -81 | 18 |
|  |  | Insular | 54 | -30 | 27 | 9 |
|  |  | RO | 52 | -51 | -18 | 15 |
|  |  | IFG | 47 | -36 | 6 | 24 |
|  |  | SMG | 46 | 6 | 63 | 18 |
|  | Main effect of TMS | SPL | 229 | 24 | -72 | 48 |
|  |  | MOG |  | 33 | -69 | 33 |
|  |  | Precuneus |  | 12 | -57 | 60 |
|  |  | Supramarginal gyrus | 116 | -45 | -36 | 24 |
|  |  | STG |  | -57 | -45 | 15 |
|  |  | Precuneus | 96 | -9 | -57 | 60 |
|  |  | SPL |  | -21 | -54 | 54 |
|  |  | MCC | 34 | 0 | -36 | 51 |
|  | Interaction Task x Site | S1 | 155 | 48 | -27 | 48 |
|  |  | S1 | 123 | -36 | -30 | 54 |
|  |  | IPL |  | -57 | -27 | 45 |
|  |  | M1 | 122 | 36 | -15 | 51 |
|  |  | SFG |  | 27 | -9 | 66 |
|  |  | M1 | 91 | -33 | -15 | 54 |
| Rest M1 stimulation | TMS > NoTMS | STG/SII | 76 | -39 | -39 | 18 |
|  | NoTMS > TMS | SOG | 48 | -21 | -72 | 39 |
|  |  | Precuenus | 31 | 3 | -48 | 48 |
|  |  | SOG | 30 | 27 | -69 | 30 |
| Vertex stimulation | TMS > NoTMS | ─ | ─ | ─ | ─ | ─ |
|  | NoTMS > TMS | MOG | 768 | -30 | -84 | 24 |
|  |  | SOG |  | 30 | -75 | 42 |
|  |  | Precuenus |  | 39 | -66 | 27 |
|  |  | SFG | 157 | -21 | 0 | 48 |
|  |  | MFG |  | -30 | 6 | 57 |
|  |  | mPFC | 107 | 9 | 54 | -9 |
|  |  | MFG | 102 | 27 | 27 | 39 |
|  |  | Precuenus | 21 | 0 | -63 | 51 |

Table S2. The results of GLM analysis

Supplementary Table 3

| Component | Cluster region | Cluster extent | Peak MNI coordinate | | |
| --- | --- | --- | --- | --- | --- |
|  |  |  | x | y | z |
| C01 (L.MN) | M1 | 1843 | -30 | -24 | 60 |
|  | S1 |  | -54 | -18 | 45 |
|  | MCC |  | -6 | -15 | 45 |
|  | STG | 438 | -45 | -27 | 12 |
|  | RO |  | -51 | -18 | 15 |
|  | Insular |  | -33 | -9 | 12 |
|  | Putamen |  | -27 | -3 | -3 |
|  | Thalamus |  | -12 | -30 | 3 |
| C02 (R.MN) | M1 | 3557 | 51 | -9 | 45 |
|  | S1 |  | 27 | -36 | 63 |
|  | SMA |  | 9 | -21 | 57 |
|  | MCC |  | 9 | 3 | 45 |
|  | RO |  | 39 | -21 | 20 |
|  | Thalamus |  | 27 | -36 | 63 |
| C04 (aDMN) | mPFC | 4420 | -3 | 48 | -6 |
|  | ACC |  | -6 | 45 | -3 |
|  | SMG |  | 12 | 60 | 9 |
|  | Caudate |  | 18 | 24 | -3 |
| C05 (R.FPN) | MFG | 4286 | 45 | 18 | 45 |
|  | IFG |  | 51 | 27 | 18 |
|  | IPL | 1386 | 51 | -51 | 42 |
|  | AG |  | 33 | -57 | 45 |
|  | Supramaginal gyrus | | 57 | -39 | 45 |
|  | MTG |  | 66 | -42 | 6 |
| C06 (L.FPN) | IFG | 5534 | -42 | 12 | 24 |
|  | AG |  | -36 | -63 | 42 |
|  | IPL |  | -33 | -54 | 42 |
| C07 (SN) | ACC | 2339 | 0 | 36 | 24 |
|  | MCC |  | 3 | 21 | 36 |
|  | Caudate | 333 | 12 | 6 | 9 |
|  | Insular |  | 36 | 15 | 0 |
|  | Putamen |  | 24 | 9 | 3 |
|  | Thalamus |  | 9 | -21 | 0 |
|  | Caudate | 296 | -12 | 9 | 6 |
|  | Insular |  | -42 | 12 | -6 |
|  | Putamen |  | -27 | 6 | 3 |
| C08 (VN) | Calcarine gyrus | 2208 | 18 | -63 | 15 |
|  | Cuenus |  | 21 | -75 | 21 |
|  | MOG |  | 36 | -78 | 15 |
| C09 (vl.FPN) | IFG | 2502 | -45 | 24 | -12 |
|  | SMG |  | -3 | 48 | 45 |
|  | AG | 645 | -57 | -57 | 33 |
|  | Supramaginal gyrus | | -51 | -66 | 36 |
|  | MTG |  | -54 | -60 | 21 |
|  | IFG | 188 | 51 | 21 | -9 |
| C10 (DAN) | SPL | 2827 | -18 | -63 | 57 |
|  | Precuenus |  | -6 | -66 | 54 |
|  | SFG |  | -18 | -1 | 60 |
|  | SFG |  | 36 | 4 | 60 |
|  | IPL | 172 | -54 | -42 | 48 |
| C11 (RON) | STG | 2687 | 66 | -24 | 12 |
|  | Supramaginal gyrus | | 66 | -21 | 18 |
|  | RO |  | 63 | 6 | 15 |
|  | STG | 2497 | -57 | -9 | 6 |
|  | Supramaginal gyrus | | -63 | -27 | 24 |
|  | RO |  | -60 | 0 | 12 |
| C12 (IN) | Insular | 5343 | -39 | 6 | -3 |
|  | Insular |  | 42 | 15 | -6 |
| C13 (pDMN) | Precuneus | 2089 | 0 | -54 | 36 |
|  | AG | 555 | -42 | -57 | 45 |
|  | AG | 454 | 54 | -51 | 27 |
| C14 (DMN) | Precuneus | 1263 | -6 | -57 | 9 |
|  | PCC |  | -6 | -51 | 24 |
|  | mPFC | 859 | 3 | 60 | -3 |
|  | SMG |  | 0 | 63 | 6 |
|  | AG | 348 | -48 | -69 | 24 |
|  | AG | 195 | 48 | -63 | 24 |

Table S3. 13 networks estimated from ICA

Supplementary Table 4

| Network | Network | F | p |
| --- | --- | --- | --- |
| C01 (L.MN) | C02 (R.MN) | 10.218 | < 0.001 |
| C02 (R.MN) | C14 (DMN) | 3.17 | 0.05 |
| C04 (aDMN) | C09 (vL.FPN) | 4.189 | 0.024 |
| C04 (aDMN) | C12 (IN) | 6.313 | 0.005 |
| C08 (VN) | C12 (IN) | 3.547 | 0.04 |
| C08 (VN) | C14 (DMN) | 5.261 | 0.01 |
| C09 (vL.FPN) | C10 (DAN) | 3.588 | 0.039 |
| C10 (DAN) | C14 (DMN) | 3.179 | 0.05 |
| C11 (RON) | C14 (DMN) | 3.383 | 0.046 |
| C12 (IN) | C14 (DMN) | 7.964 | 0.002 |

Table S4. The group effect in the FNC between networks
